# Testing various combinations of cryoprotective agents for human skin cryopreservation by vitrification method

**DOI:** 10.1101/2024.12.21.629872

**Authors:** Andrei Riabinin, Maria Pankratova, Olga Rogovaya, Ekaterina Vorotelyak, Andrey Vasiliev

## Abstract

Modern translational medicine, pharmacology and other fields require effective skin cryopreservation technology. Vitrification is promising technology for this aim. This rapid freezing technique minimizes hexagonal ice crystals formation, time of stress condition for cells in samples and dehydration both in the cytoplasm of cells and in the extracellular space. This study is aimed at finding the optimal combination of extracellular (sucrose, mannitol, polyethylene glycol and FBS) and intracellular (DMSO, glycerol, ethylene glycol) cryoprotective agents to achieve the highest viability and tissue structure preservation of skin samples after vitrification.

## Introduction

Cryopreservation is a process during which biological material is cooled to sub-zero temperatures to be stored for a long period of time while completely stopping metabolism and preserving the viability of the samples after they are frozen (Hunt, 2017). One of the main biological objects for which various cryopreservation techniques can be effectively applied and many freezing protocols have been developed is the human skin, skin components and living equivalents of skin components. In addition, the development of cryopreservation technology is essential for several scientific fields, particularly translational medicine and cell biology.

Current progress in optimizing cryopreservation approaches aims to increase cell viability after undergoing a full cryopreservation cycle, to minimize damage to extracellular structures of tissues and organs, and to optimize and reduce the cost of the process itself (Chen et al., 2023). One of the important aspects required for effective tissue cryopreservation is the selection of cryoprotective agents (CPAs). The protective effect of CPAs is based on their ability to influence the water crystallization process, reduce the number of ice crystals formed and increase the percentage of water in the liquid or amorphous ice phase at sub-zero temperatures, which minimizes physical effects on cellular and extracellular matrix components, preserve proteins structure and to stabilize cell membrane (Bojic et al., 2021; Whaley et al., 2021, Chang et al, 2005, Patel et al, 2023). CPAs are cytotoxic in most cases and have varying rates of penetration and efficacy within the preservation of different biological samples, which significantly limits their maximum concentration and applicability to specific tasks and biological targets (Whaley et al., 2021). Skin cryopreservation often uses a combination of intracellular and extracellular CPAs, but the composition of the cryopreservation medium and the concentration of CPAs varies considerably between protocols (Bravo et al., 2000; Erdag et al., 2002; Kua et al., 2012; Schiozer et al., 2013; Xu et al., 2022).

Another key factor ensuring successful cryopreservation of samples is the choice of the optimal freezing temperature regime. The two main approaches within this aspect are systemic slow freezing and vitrification – instantaneous freezing whereby the liquid within the sample enters an amorphous ice state at -130°C, by passing the crystallization phase with the formation of ice crystals (Fahy et al., 2015; Whaley et al., 2021). According to studies in which the vitrification method has been applied to human skin, this method of cryopreservation is quite effective and demonstrates an identical or higher level of preservation of sample integrity and cell viability in its composition compared to slow freezing (Fujita et al., 2000; Campbell et al., 2022). In addition, tissue viability is influenced by the duration and temperature conditions of sample storage. A number of studies have shown that skin samples subject to long-term cryopreservation should preferably be stored at temperatures below -130°C in vapour nitrogen or at -196°C in liquid nitrogen to avoid ice crystal formation and cell damage (Udoh et al., 2000; Schiozer et al., 2013; Pianigiani et al., 2016). At sufficiently low storage temperatures, allografts remain viable even after long-term storage (Udoh et al., 2000; Ben-Bassat et al., 2001).

An important challenge in skin cryopreservation is to ensure the long-term viability of the specimen (Kagan et al, 2005). The results of viability assessment after long-term skin cryopreservation suggest that under low-temperature storage conditions (below -130°C), samples remain viable for up to 5 years (Bravo et al., 2000; Schiozer et al., 2013). While the properties of CPAs such as glycerol, ethylene glycol, dimethyl sulfoxide (DMSO) and fetal bovine serum (FBS) are well-tested (Mandumpal et al, 2011; Eriani et al, 2021; Kang et al, 2012; May et al, 1980; Fujita et al, 2000; Teasdale et al, 1993), the cryoprotective properties of substances such as polyethylene glycol (PEG) and mannitol have not been well studied and have not been tested for skin vitrification. Mannitol is a sugar and can form amorphous glass during vitrification. Two mechanisms have been proposed for amorphous sugars to provide cryoprotection. The first suggests that the proteins are mechanically immobilized in the glassy matrix and the restriction of motion and relaxation processes is thought to prevent protein unfolding (Chang et al, 2005). The second mechanism is referred to as the ‘water replacement’ or ‘preferential exclusion’ hypothesis (Arakawa et al, 1985; Carpenter et al, 1988). According to this theory, sugars protect proteins by hydrogen bonding to the exposed polar and charged group on the surface of proteins. It was shown that mannitol supplementation improved the efficiency of follicles cryopreservation (Kiroshka et al, 2018). PEG also stabilizes proteins in the sample but does so due to the direct formation of peptide bonds with them. In addition, PEG acts as a solubilizer and, as we assume, promotes better dissolution and, consequently, higher activity of other CPAs in the cryopreservation agents. Their efficiency as CPAs was demonstrated in mesenchymal stem cells (MSCs) cryopreservation (Patel et al, 2023). The aim of this work was to determine the optimal combination of CPAs for effective vitrification of skin and epidermal layers.

## Materials and methods

### Samples

Skin samples were obtained during plastic surgery procedures at the Medical Research Center of High Medical Technologies – Central Military Clinical Hospital named after A.A. Vishnevsky of the Ministry of Defense of the Russian Federation in accordance with the Agreement on scientific cooperation between this center and N.K. Koltzov Institute of Developmental Biology of the Russian Academy of Sciences (IDB RAS) with the informed consent of the patients.

Skin fragments and epidermal layers from 3 different donors were used in the experiments (N=3). For each experimental group, 5 technical repetitions were performed.

To construct human skin equivalents (HSEs), a matrix (hyaluronan-collagen membrane G-Derm (G-grupp Ltd.) was prepared for cell seeding as follows: G-Derm were cut out on 4×4 cm slices and placed it in a sterile Petri dish. The membrane pieces were washed once with Hanks’ solution. MSCs were seeded on the prepared membranes (cells were detached according to the standard protocol) at a concentration of 3×10^4^ cells/cm^2^. After 3 hours of incubation, keratinocytes were planted on the surface of the matrix with attached MSCs at a concentration of 1×10^5^ cells/cm2. The matrix with cells was cultured in the nutrient medium for keratinocytes at 37°C in an atmosphere of 5% CO2 until the keratinocytes formed an epithelial layer on the matrix surface.

### Processing of donor material

After receiving the skin, it was washed in sufficient volumes of Hanks solution (PanEco) with an antibiotic (gentamicin (PanEco)). Then the hypodermis (fat) was cut off using a scalpel, and the flap of skin was cut into equal round fragments with a diameter of 5 mm using a scalpel.

To obtain skin sheets, some of the resulted skin fragments were incubated for 24 hours at +4 °C in 0.2% dispase (Sigma). After that, the epithelial layer was mechanically separated from the dermis using forceps.

To disaggregate keratinocytes, epidermal fragments were placed in a tube in a solution consisting of 0.25% trypsin (Gibco) and 1:1 phosphate buffered saline (PBS) (PanEco) and incubated at 37°C for 10-15 minutes under visual inspection and periodic shaking of the tube. Trypsin action was inhibited by the addition of FBS (Capricorn scientific) at 2% of the total liquid volume. The suspension with epidermal pieces was pipetted and the resulting suspension of keratinocytes was centrifuged for 5 minutes at 300g (1500 rpm).

After centrifugation, the supernatant was drained and the cell sediment was resuspended in a mixture of DMEM/F12 media (PanEco) supplemented with 10% FBS, 10 ng/ml epidermal growth factor (EGF) (Sigma), 1% Glutamax (Gibco), 1% penicillin-streptomycin (PenStrep) (Gibco) and 1% ITS-X (100X) (Gibco) (keratinocyte culture medium). 100 µl of suspension was sampled to count the number and viability of cells. Cells were stained with 0.4% trypan blue at a 1:1 dilution and the number of viable cells was determined using a NanoEntek EVE Plus automated cell counter. The resulting suspension was seeded at a concentration of 2×105 cl/ml into cell culture flasks pre-sorbed with collagen solution. Keratinocytes were cultured at 37°C in an atmosphere of 5% CO2.

After one day, fresh keratinocyte culture medium with 5mM Rock-inhibitor (STEMCELL Technologies) was added to the flasks, and then the culture medium was changed every 2-3 days. After the cells filled 70-80% of the area of the culture flasks, they were removed from the surface by treatment with 0.25% trypsin, resuspended and dispersed into new flasks at a ratio of 1:3.

Homogenized fat was transferred into 50 ml tubes, 27 ml of DMEM/F12 medium was added to each tube, and one aliquot of each type of collagenase (1.5 ml of 2% collagenase I and II solution (Wohrtinghton)) was added. The tubes were inverted to mix the contents, placed in a thermostat and incubated for no more than 2 hours at 37°C with constant stirring on an orbital shaker. After incubation, 1 ml of FBS was added to each tube and centrifuged (300g, 10 min) at 4°C. The supernatant and fat layer were carefully withdrawn, 10-15 ml of DMEM/F12 medium were added and centrifuged (300g, 10 min).

The supernatant was removed, erythrocyte lysis buffer was added to each tube (9 ml of buffer per 1 ml of cell sediment), incubated for 10 min at room temperature and centrifuged (300g, 10 min). The cell sediment was resuspended in 10 ml of MSC culture medium (DMEM/F12, 10% FBS, 1% Glutamax, 1% ITS-X (100X), 1% PenStrep) and centrifuged (300g, 10 min). The supernatant was removed, and the precipitate was resuspended in 10 ml of culture medium. 100 µl of suspension was taken for cell counting and viability (using the above method). From each 50 ml tube, the cell suspension was dispensed into one flask with a surface area of 75 cm^2^ (T75) and cultured in an incubator at 37°C in an atmosphere of 5% CO2. After 24 hours, the number of attached cells was evaluated, and if the confluency was at least 40%, the suspension with tissue fragments (if available) was transferred to a new flask, and 12 ml of culture medium was added to the old flask. The medium was changed every 2-3 days until 80-90% of the surface area of the culture flask was covered with cells.

The isolated donor cells were screened for phenotypic markers of MSCs (expression of CD90, CD44, CD29, CD73 and CD105, and no expression of CD34, CD45, CD31).

### Skin cryopreservation

5×5 mm skin layers and fragments were incubated for 30 minutes in the cryopreservation medium (Table 1) in sterile retort bags at room temperature. Three skin fragments and three epithelial layers were placed in each bag. After incubation, the bags were sealed, placed in liquid nitrogen and transferred to the cryopreservation room.

**Table 1.** Compositions of cryopreservation media.

| **Cryopreservation medium (Glycerol + FBS)** |  |  |
| --- | --- | --- |
| Component | Manufacturer | Concentration |
| RPMI-1640 | PanEco | 1x |
| Glycerol | Sigma | 20% |
| FBS | Capricorn | 10% |
| EGF | PanEco | 10 ng/ml |

| **Cryopreservation medium (DMSO + PEG + sucrose)** |  |  |
| --- | --- | --- |
| RPMI-1640 | PanEco | 1x |
| DMSO | Sigma | 20% |
| Polyethylene glycol 6000 | Sigma | 20% |
| Sucrose | Sigma | 0.5M |
| Epidermal growth factor | PanEco | 10 ng/ml |
| **Cryopreservation medium 3 (Glycerol + FBS + mannitol)** |  |  |
| RPMI medium | PanEco | 1x |
| Glycerol | Sigma | 20% |
| FGF | Capricorn | 10% |
| Mannitol | Macklin | 0.5 M |
| Epidermal growth factor | PanEco | 10 ng/ml |
| **Cryopreservation medium 4 (Ethylene glycol (EG) + PEG + sucrose)** |  |  |
| RPMI-1640 | PanEco | 1x |
| Ethylene glycol | Sigma | 20% |
| Polyethylene glycol 6000 | Sigma | 20% |
| Sucrose | Sigma | 0.5M |
| EGF | PanEco | 10 ng/ml |

### Cryopreservation medium cytotoxicity testing

Fragments of control skin samples were transferred into 48-well plates before freezing and incubated for 30 min at 37°C. After 30 minutes incubation, MTT or resazurin assay was performed according to the protocols described below.

### Thawing of samples

Defrosting of the sample bags was performed one week after cryopreservation by immersing them in a water bath at 37°C for 1 minute until the contents of the cryopack were completely converted to a liquid state. 10 ml of RPMI-1640 medium (PanEco) was added to thawed samples, to gradually reduce the osmotic pressure of the solution. Then the samples were washed once from CPAs in PBS and transferred to the wells of a 48-well plate (Fig. 1)

**Figure 1.**
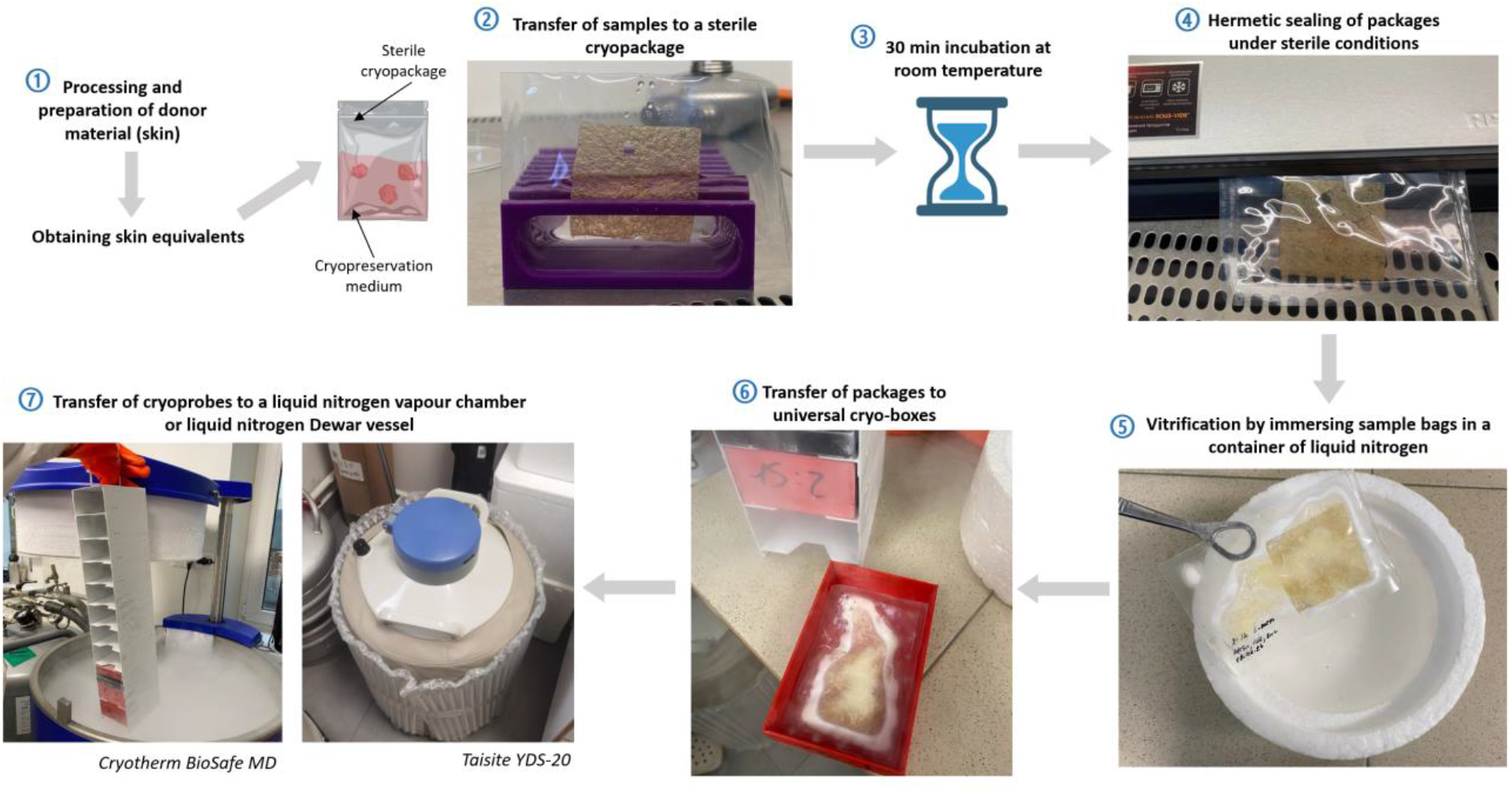
Cryopreservation protocol.

### Measuring the viability of samples after defrosting

#### MTT test

300 µl of RPMI medium with 10% MTT solution (PanEco) was added to each well of a 48-well plate containing control and thawed samples. The plate was kept in the incubator for 4 hours at 37°C. The MTT medium was then withdrawn from the wells and 300 µl of DMSO was added to each well. Dissolution of the resulting formazan in DMSO was carried out on a 3d-shaker at 60 rpm for 12 hours. After the appropriate incubation time, 100 µl of solution was transferred into the wells of a 96-well plate. The plate was analyzed on a BioTek Synergy H1 spectrophotometer at 545 nm (Fig. 1).

Skin and epithelial samples treated with a combination of detergent solutions (10% Triton X-100 (Sigma-Aldrich), 10% sodium dodecyl sulfate (Sigma-Aldrich) and 10% TWEEN 20 (AppliChem) in PBS) for 30 minutes at 37°C served as negative controls. Unfrozen samples were used as a positive control. To standardize the results of formazan concentration measurements by weight of the original samples, the values obtained on the spectrophotometer were divided by the weight of the corresponding skin fragments.

#### Resazurin test

300 µl of 1% resazurin solution (Abisense) was added to each well of a 48-well plate containing control and thawed samples. The plate was incubated for 4 hours in an incubator at 37°C and then 100 µl of solution was transferred to each well of a 96-well plate. The plate was analyzed on a BioTek Synergy H1 spectrophotometer at an excitation wavelength of 570 nm and emission wavelength of 600 nm (Figure 2). Skin and epithelial layer samples treated with combinations of detergent solutions (10% Triton X-100, 10% sodium dodecyl sulfate and 10% TWEEN 20 in PBS) for 30 minutes at 37°C served as negative controls. Unfrozen samples were used as a positive control. To standardize the results of formazan concentration measurements by the weight of initial samples, the values obtained on the spectrophotometer were divided by the weight of the corresponding skin fragments.

**Figure 2.**
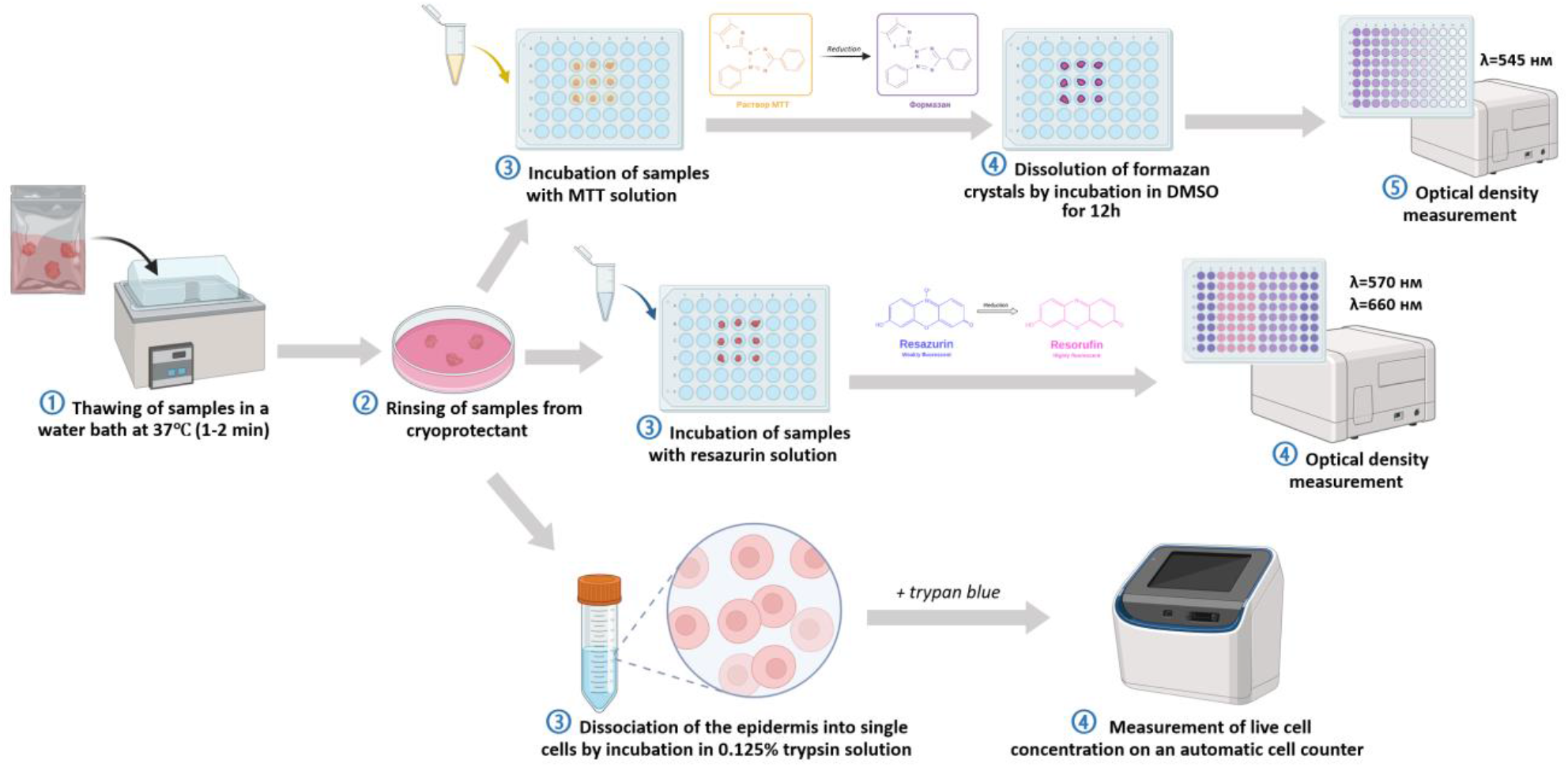
Schematic for thawing samples and assessing their viability: performing MTT and resazurin tests, staining keratinocytes isolated from epidermal layers with trypan blue.

#### Assessment of epidermal layer viability using trypan blue

Control and thawed epidermal strata were incubated for 20 min in 0.125% trypsin (PanEco) at 37 °C. An equal volume of PBS with 10% FBS was added to the obtained suspension and centrifuged (250g, 5 min). The supernatant was removed and 1000 µl of RPMI medium was added to the precipitate and pipetted. Cells were stained with 0.4% trypan blue at a dilution of 1:1 and the number of viable cells was determined using an automatic cell counter NanoEntek EVE Plus (Fig. 2).

#### Histological staining

For histological analysis, skin and epidermal fragment samples were cut into 5×5 mm fragments, fixed in 10% formalin for 24 hours, dehydrated, embedded in paraffin and 5 µm thick sections were obtained. For haematoxylin-eosin staining, dried skin sections were stained with haematoxylin for 5 minutes after standard histological wiring through alcohols of descending strength, washed in running water for 10 minutes and dipped in distilled water for 2-3 seconds. Eosin was applied for 7 seconds and rinsed with distilled water.

For Van Gieson staining, dried glass with sections after standard histological wiring through alcohols of descending strength was placed in distilled water, iron haematoxylin was applied according to Weigert for 10 minutes, washed with tap water for 10 minutes. The slices were then covered with Van Gieson’s picrofuchsin for 10 minutes and dipped in distilled water for 2-3 seconds.

After staining, slices were run in alcohols of increasing strength and enclosed under cover glass in mounting medium Vitrogel (Biovitrum). Histological preparations were visualized on an Olympus IX73 inverted fluorescence microscope with an Olympus DP74 camera (Olympus).

#### Statistical analysis

All samples were checked for normal distribution using the Shapiro-Wilk test in the JAMOVI programme (v.2.3.28). Results with p values > 0.05 supported the null hypothesis of normality of distribution. Statistical analysis of the obtained results of sample viability was performed using STATISTIKA software (v.10.0). Significant differences between the two groups (negative control and experimental groups) were examined using one-factor analysis of variance (ANOVA). Data in the graphs are presented as mean ± standard deviation. The critical level of significance was assumed to be 0.05.

## Results and discussion

### Study of cytotoxicity of cryopreservation medium

Since the cryopreservation protocol developed by us includes incubation of fresh samples in cryopreservation medium for 30 minutes, we tested the cytotoxicity of the most effective cryopreservation medium based on the tests performed in the previous stage of work on two types of samples: human skin and HSEs. The viability of skin fragments incubated in the cryopreservation medium with glycerol and FBS was not different from the samples in RPMI medium without added CPA (Fig. 3A). Similar results were observed for the combination of DMSO, PEG and sucrose (Fig. 3A). For HSEs, MTT analysis also showed no significant decrease in viability after incubation in both formulations tested (Figure 3B).

**Figure 3.**
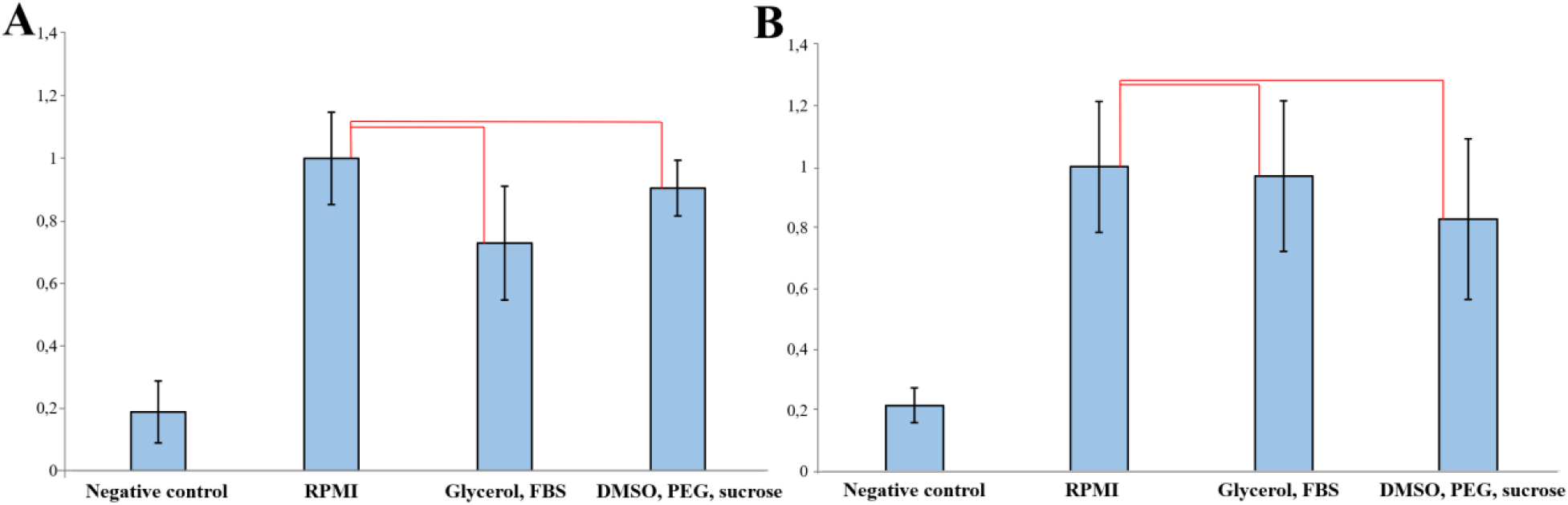
Evaluation of cytotoxicity of the cryopreservation medium 1 and 2. Relative light absorption coefficients of formazan solutions in skin samples (A), HSE (B) after 30 minutes of incubation of tested cryopreservation medium. Scale: 1 is 100% of cell viability (positive control).* - statistically significant difference; p<0.05.

The data obtained allow us to conclude that there is no detectable adverse effect of the cryopreservation media used on the viability of skin and its combined biological equivalent.

### Assessment of skin and epidermal strata viability after vitrification and storage for one month

Our previous work showed that vitrification is an effective way to freeze fragments of human skin and epidermal strata that allows us to preserve cell viability at a high level using several combinations of CPAs (Riabinin et al., 2023). The most effective of the tested CPAs for vitrification and short-term (one week) storage of skin and epidermis was a combination of 20% glycerol and 10% FBS, as well as a cryopreservation medium including 20% DMSO, 20% PEG and 0.5M sucrose (Riabinin et al., 2023).

In the current work, both cryopreservation media have been tested for vitrification of skin and epidermis samples. In addition, since one of the key objectives of cryopreservation within our subject is long-term storage of viable samples for further use for transplantation to patients, the storage time of the samples was increased to one month.

The results obtained when assessing the viability of the specimens after thawing using MTT assay showed that cryopreservation medium based on glycerol and FBS, and those based on a combination of DMSO, PEG and sucrose could preserve the viability of skin fragments at a high level, Δ(MTT value) for these groups was -18% and -20%, respectively (Figure 4A).

**Figure 4.**
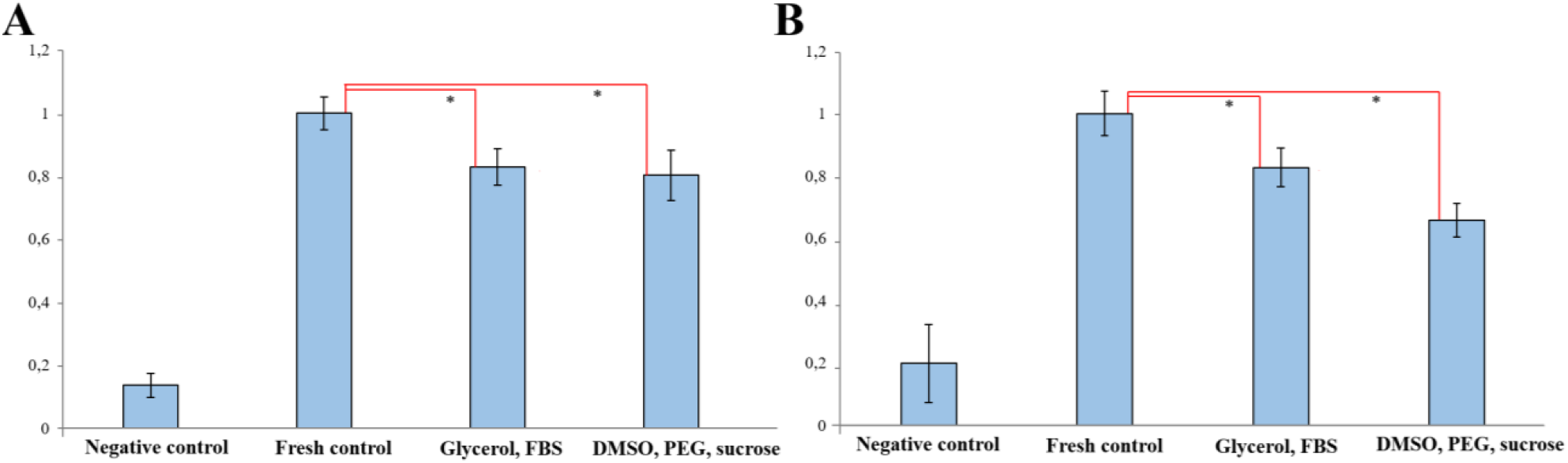
Assessment of viability of skin (A) and epithelial layers (B) samples. Relative light absorption coefficients of formazan solutions in samples after freezing for one month. Scale: 1 is 100% of cell viability (positive control). * - statistically significant difference; p<0.05.

In the case of epidermal strata, the cryopreservation medium with the addition of glycerol and FBS was more effective (Δ(MTT value) = -18%) compared to cryopreservation medium 2 (Δ(MTT value) = -35%), which is broadly consistent with our previous results obtained with shorter sample storage times (Figure 4B) (Riabinin et al., 2023).

Live cell counting was also performed in the suspension of keratinocytes isolated from epithelial layers. The survival rate in the control group was 45.6%. After freezing, the percentage of live cells was 27.3% in the group with glycerol-serum-based cryoprotectant, and for the combination of DMSO, PEG and sucrose -21.1% (Fig. 5). These results are not fully consistent with the results of viability assessment by MTT assay, which is probably due to the nature of this technique. In particular, a significant decrease in cell survival could be due to prolonged enzymatic treatment of epithelial layers during sample preparation. In addition, due to the limited amount of cellular material, standard deviations and statistics are not presented in the graphs.

**Figure 5.**
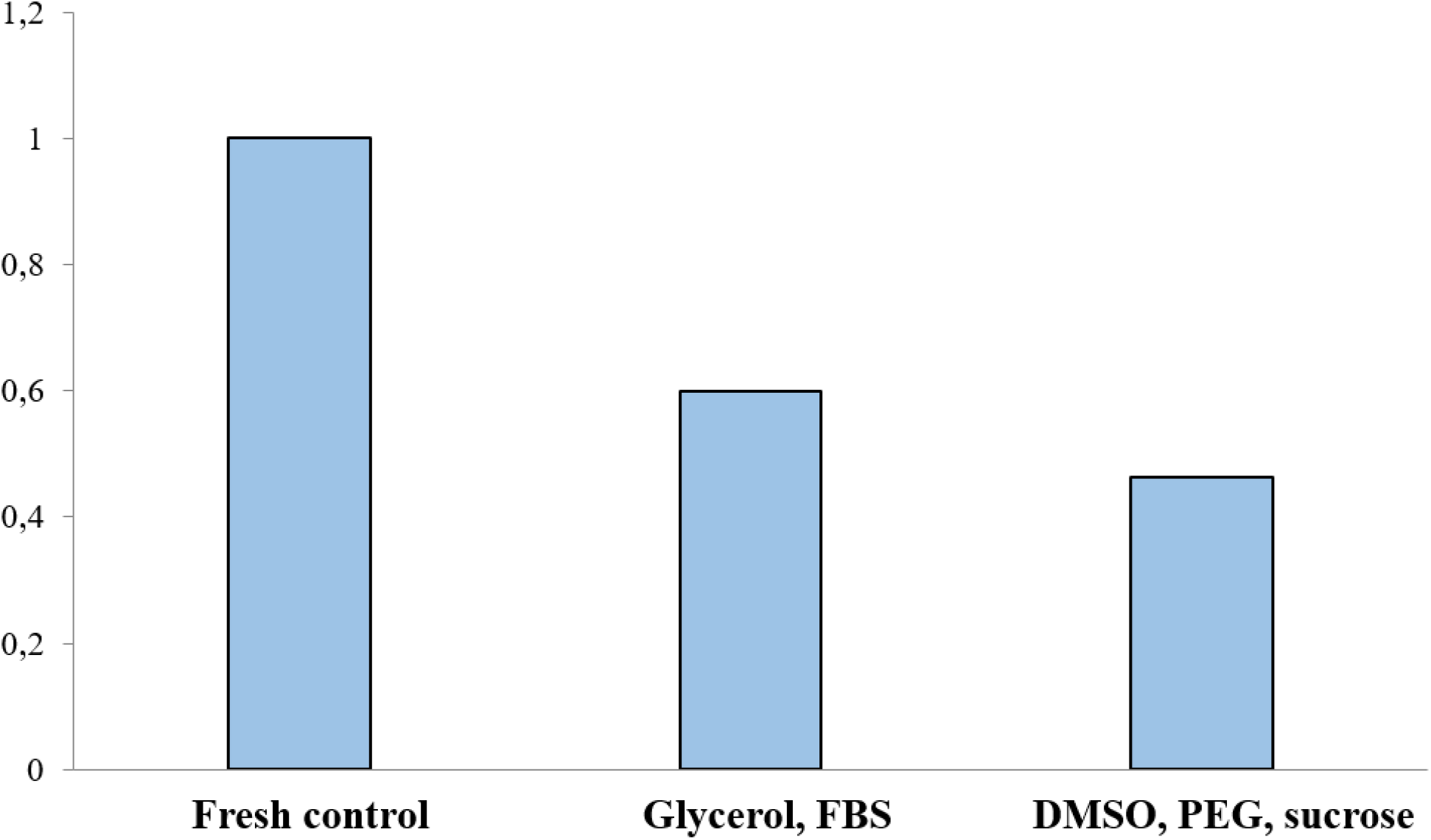
Concentration of live cells after trypan blue staining of keratinocyte suspension isolated from epithelial layers. Scale: 1 is 100% of cell viability (positive control).

Thus, we found no noticeable decrease in skin and epidermal viability when the sample storage time was increased to one month. These results demonstrate that the selected cryopreservation medium maintains high level of tissue viability both during short-term and longer cryopreservation.

### Assessment of HSEs viability after storage in different temperature regimes

Cryopreservation of skin equivalents and the development of a storage system for cryopreserved specimens are critical for their effective use in clinical practice. Proper cryopreservation techniques allow equivalents to be stored for weeks or months while still being available for immediate use. According to the literature, vitrification of skin equivalents has only recently been tested and, despite the potential for application and further development, is currently poorly understood, particularly due to the diversity of equivalent types (Campbell and Brockbank, 2022).

At this stage of our work, the efficiency of the selected cryopreservation medium in vitrification of HSEs was also tested. In addition, two different temperature regimes of sample storage were tested: at around -130°C in vapour nitrogen or at -196°C in liquid nitrogen. The results of MTT analysis of the samples after thawing showed that the combination of glycerol and FBS maintained the viability of HSEs at a high level after storage at both -130°C (Δ(MTT value) = -24.7%) and -196°C (Δ(MTT value) = -23.6%) (Figure 6). Similar viability values were obtained for the cryopreservation medium with the addition of DMSO, PEG and sucrose: Δ(MTT value) after cryopreservation at -130°C was -24.1%, and for -196°C this value was -24.5%.

**Figure 6.**
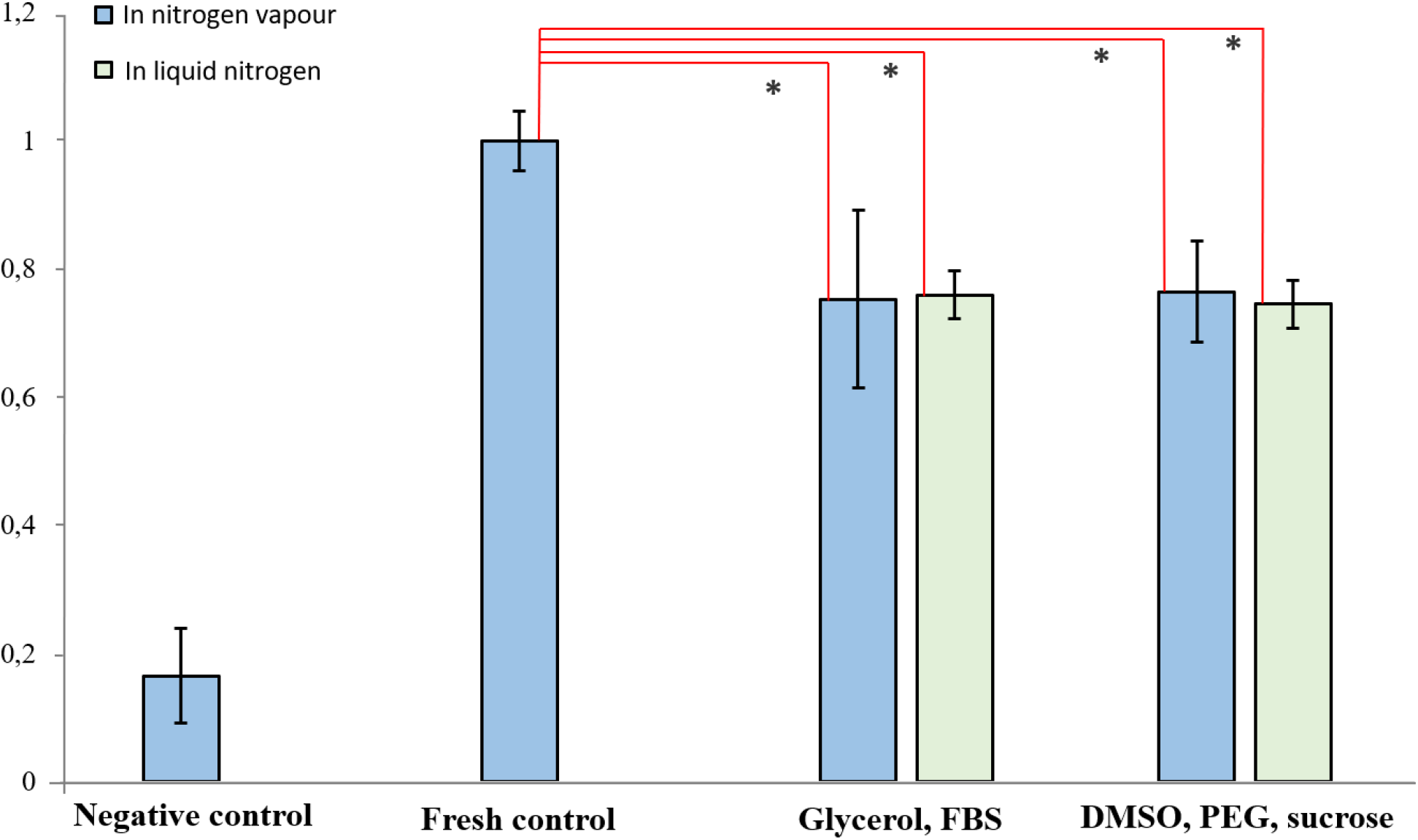
Assessment of HSEs viability. Relative light absorption coefficients of formazan solutions in negative control, control before cryopreservation and experimental groups after one month of cryopreservation in vapour and liquid nitrogen. Scale: 1 is 100% of cell viability (positive control).* - statistically significant difference; p<0.05.

In addition, in the course of this work cryopreservation medium based on mannitol and ethylene hydrochloride was tested.

First, the cytotoxicity of the developed cryopreservation medium was evaluated. The viability of fresh skin fragments incubated in the cryopreservation medium based on ET, PEG and sucrose did not differ from the control samples in RPMI medium without added CPA. At the same time, for the combination of glycerol, FBS and mannitol, the resazurin test revealed a significant decrease in cell viability in the samples compared to control, which indicates a cytotoxic effect of this cryopreservation medium (Fig. 7).

**Figure 7.**
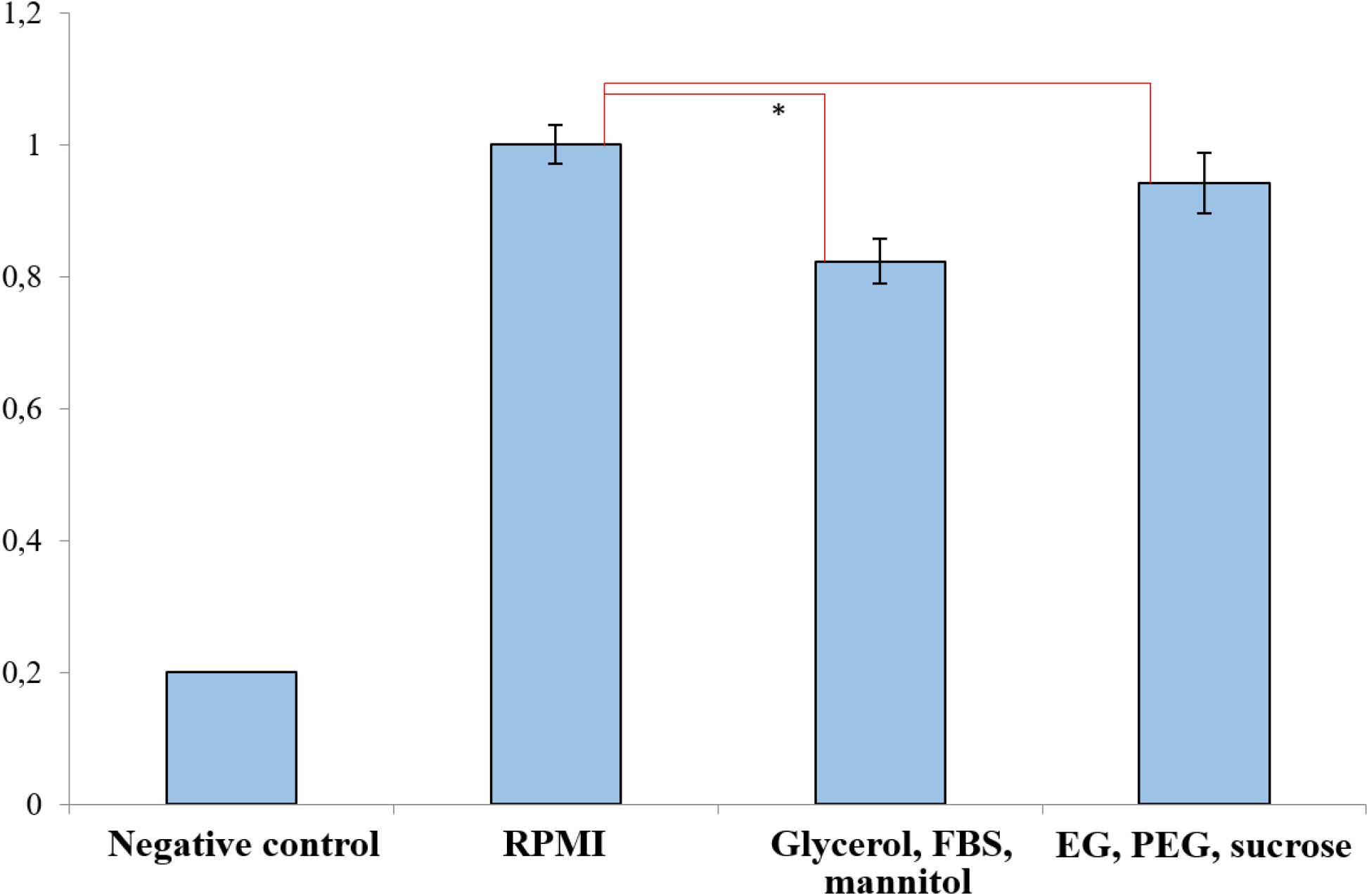
Evaluation of cytotoxicity of the developed cryopreservation medium. Relative light absorption coefficients of resorufin solutions in skin samples after 30 minutes of incubation of tested cryopreservation medium. Scale: 1 is 100% of cell viability (positive control). * - statistically significant difference; p<0.05.

At this stage of the work, the resazurin test was used to assess cell viability of the samples, instead of the previously used MTT test. According to literature data, both tests are suitable for analyzing skin viability (Cook et al., 2007; Landsman et al., 2016). Like the MTT, the resazurin test serves to measure the metabolic activity of cells and monitor cell viability, and is based on the reduction of resazurin to pink-colored and highly fluorescent water-soluble resorufin (Longhin et al., 2022). However, unlike MTT, the resazurin test lacks the dissolution stage of formazan crystals, which significantly reduces the assay time and allows viability assessment within one day. In addition, the resazurin test has several other advantages. Resazurin is non-toxic, whereas MTT, according to some literature reports, can have cytotoxic effects when incubated with the solution for long periods of time. In addition, the resazurin test is more sensitive in assessing cytotoxicity than MTT (Ghasemi et al., 2021; Longhin et al., 2022).

Next, we evaluated the effectiveness of the developed cryopreservation media 3 and 4 for freezing skin and epidermis samples for one month. The results obtained when assessing skin viability after thawing using the resazurin test showed that the most effective of the cryopreservation medium we tested was a combination of EG, PEG, and sucrose (Δ(rezofurin value)= -20.6%). At the same time, samples frozen in a preservative based on glycerol, FBS and mannitol showed a stronger decrease in viability, Δ(rezofurin value) for this group was -41.54% (Fig. 8A). However, the differences between the relative viability of skin samples in these two experimental groups and the control group were not statistically significant.

**Figure 8.**
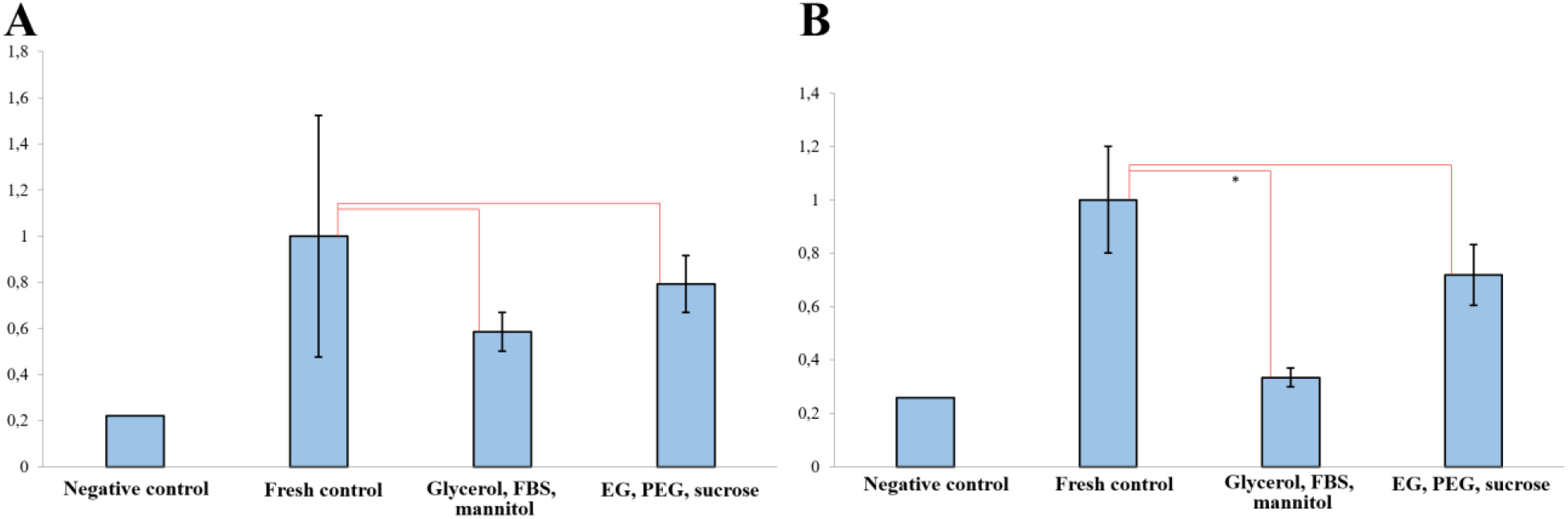
Assessment of viability of skin samples (A) and epithelial layers (B). Relative light absorption coefficients of resofurin solutions in samples after freezing for one month. * - statistically significant difference; p<0.05.

Similar trends were observed for epidermal layers. Cryopreservation medium with the addition of glycerol, FBS and mannitol was less effective, as we observed a significant decrease in cell viability (Δ(rezofurin value)= -66.56%) compared to fresh samples. The cryopreservation medium 4 allowed us to maintain cell viability at a higher level (Δ(rezofurin value = -28, 19%), no significant differences were observed with the control (Figure 8B).

Additionally, live cells were counted in a suspension of keratinocytes isolated from epithelial layers using trypan blue staining. In the group with cryopreservation medium with the addition of glycerol, FBS and mannitol, a 31% decrease in live cells was observed, while for the combination of EG, PEG and sucrose this figure was 25.3% (Fig. 9). These results are in general agreement with the results of viability assessment using the resazurin test.

**Figure 9.**
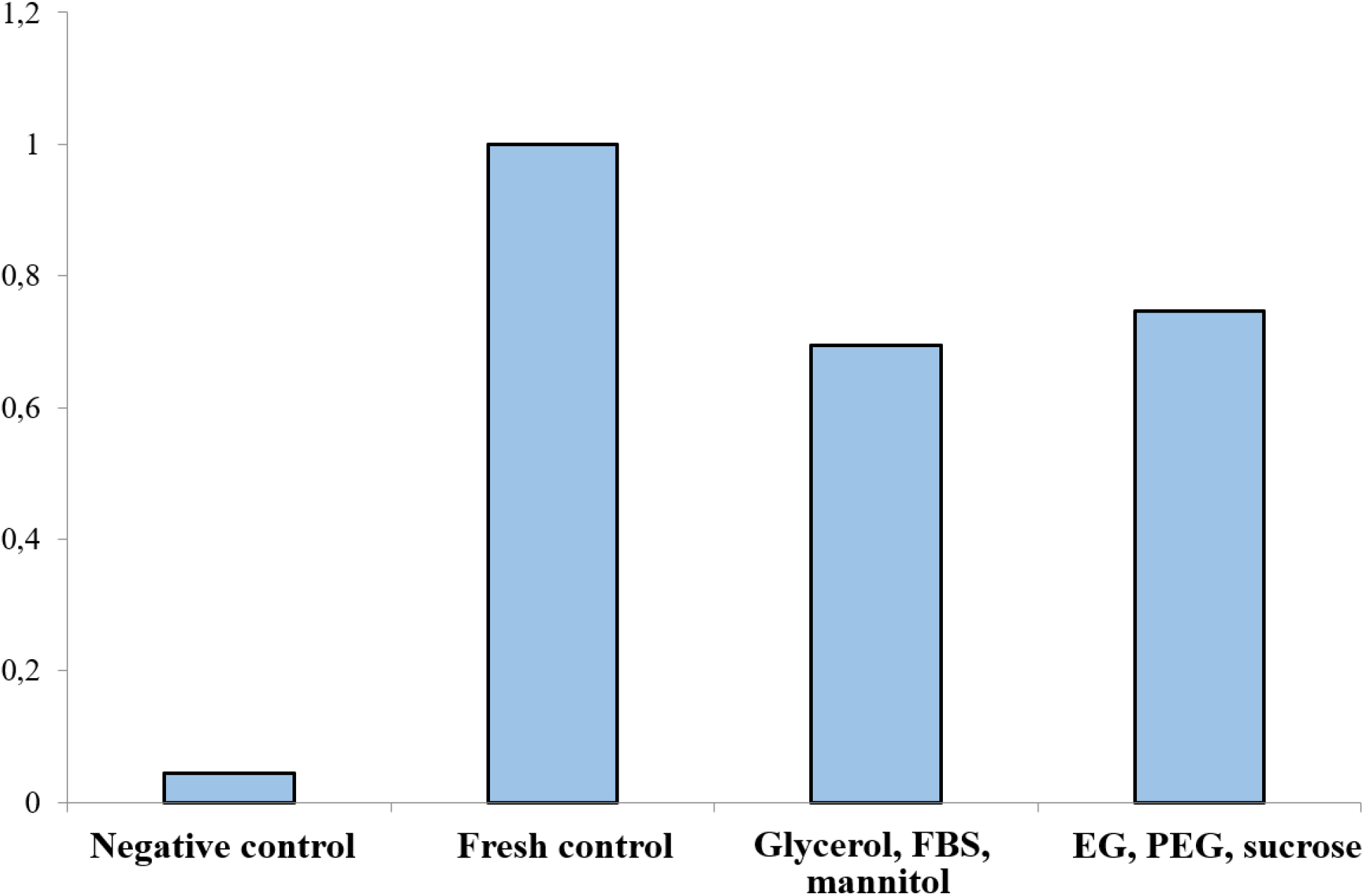
Concentration of live cells after trypan blue staining of keratinocyte suspension isolated from epithelial layers.

Thus, we found a decrease in skin and epidermis viability when samples were frozen in cryopreservation medium with the addition of glycerol, FBS, and mannitol. Given that the combinations of glycerol and mannitol, FBS allowed the viability of the samples to be maintained at high levels (Figures 3-5), we hypothesize that the addition of mannitol to the medium in combination with FBS may have created high osmotic pressure in the extracellular space, and consequently cell damage and death. At the same time, the second tested preservative based on EG, PEG and sucrose allowed to maintain high viability of both epidermis and skin.

Since effective cryopreservation aims to preserve not only the viability of cells but also the three-dimensional structure of the tissue, a histological study of cross sections of skin samples after freezing was performed (Fig. 10). H-E staining showed that in all samples, both in the dermis and epidermis, discernible cell nuclei are detected, which indicates that cryopreservation in the tested media does not destroy them. The structure of the epidermis also remained intact in both groups, but in the samples frozen in medium with glycerol, FBS and mannitol, separation of the epidermis from the dermis was observed, which is probably due to mechanical damage caused by basal membrane or adhesion contact damage during the cryopreservation cycle (Fig. 10A).

**Figure 10.**
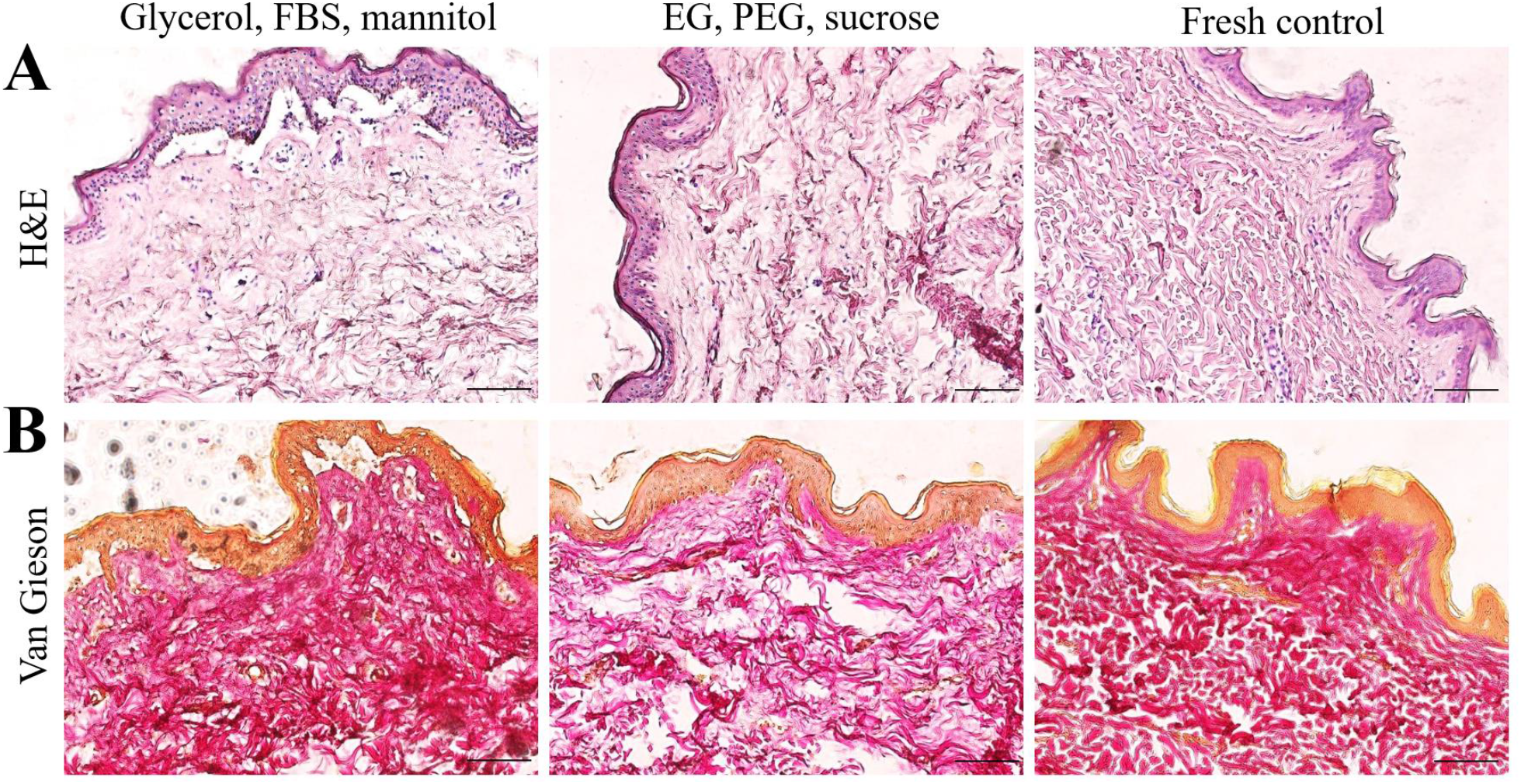
Histological analysis of skin slices after freezing. Haematoxylin-eosin (A) and Van Gieson (B) staining. Scale bar: 100 μm.

The structure of dermal collagen fibers, detected by Van Gieson staining, was altered in both samples compared to the fresh control. In the case of the cryopreservation medium with Glycerol, FBS and mannitol, a thickening of collagen fibers and a decrease in the distance between them were observed. In samples with the cryopreservation medium with EG, PEG and sucrose, the structure of the dermis was more disorganized, with a significant increase in the distance between collagen fibers in some areas (Fig. 10B). However, additional studies are required to assess the influence of the abovementioned morphological disturbances of the skin structure after freezing on the transplantation efficiency.

## Conclusions

Thus, in the current work we tested the cryopreservation medium that proved to be the most effective at the previous stage of the study (Riabinin et al, 2023) and developed two new cryopreservation media. Both cryopreservation media from the previous phase showed high residual viability even after longer (one month) cryopreservation of skin and epidermal layers.

Both cryopreservation media also provided high levels of viability during vitrification of HSEs based on hyaluronic acid hydroxycolloid and collagen with a cellular component of MSCs and primary keratinocytes. In addition, we did not detect any cytotoxic effect of the used CPAs on the viability of the skin and its combined biological equivalent. Storage regimes in liquid nitrogen (−196°C) and in liquid nitrogen vapour (−130°C) showed no significant difference in the residual viability of the samples after cryopreservation for one month.

Among all tested cryopreservation media, media based on FBS and glycerol combination and based on new combination of EG, PEG and sucrose showed highest viability of skin and epidermis after storing the samples for one month. But vitality test with trypan blue demonstrated highest viability of cells after vitrification in second one medium. Also, cryopreservation media with EG, PEG and sucrose combination and glycerol, FBS and mannitol combinations demonstrated low cytotoxicity level in comparison to previous tested. The cryopreservation medium based on glycerol, FBS and mannitol demonstrated a decrease in skin and epidermis viability, as well as disruption of epidermis and dermis structure and epidermis-dermis contact, which was confirmed by histological examination of the samples.

## Data availability

